# Analysis of an Indian colorectal cancer faecal microbiome collection demonstrates universal colorectal cancer-associated patterns, but closest correlation with other Indian cohorts

**DOI:** 10.1101/2022.10.24.513519

**Authors:** Mayilvahanan Bose, Henry M Wood, Caroline Young, International CRC Microbiome Network (AMS/CRUK), Philip Quirke, Ramakrishan Ayloor Seshadri

## Abstract

It is increasingly being recognised that changes in the gut microbiome have either a causative or associative relationship with colorectal cancer (CRC). However, most of this research has been carried out in a small number of developed countries with high CRC incidence. It is unknown if lower incidence countries such as India have similar microbial associations.

Having previously established protocols to facilitate microbiome research in regions with developing research infrastructure, we have now collected and sequenced microbial samples from a larger cohort study of 46 Indian CRC patients and 43 healthy volunteers. When comparing to previous global collections, these samples resemble other Asian samples, with relatively high levels of *Prevotella*. Predicting cancer status between cohorts shows good concordance. When compared to a previous collection of Indian CRC patients, there was similar concordance, despite different sequencing technologies between cohorts. These results show that there does seem to be a global CRC microbiome, and that some inference between studies is reasonable. However, we also demonstrate that there is definite regional variation, with more similarities between location-matched comparisons. This emphasises the importance of developing protocols and advancing infrastructure to allow as many countries as possible to contribute to microbiome studies of their own populations.

**Importance:** Colorectal cancer is increasing in many countries, thought to be partly due to the interaction between gut bacteria and changing diets. While it is known that populations in different parts of the world have very different gut microbiomes, the study of their role in colorectal cancer is almost exclusively based in the USA and Europe. We have previously shown that there is overlap between the colorectal cancer microbiome in multiple different countries, establishing robust protocols in the process. Here we expand that into a new Indian cohort. We show that while there are similarities between countries, by concentrating on one country, we can uncover important local patterns. This shows the value of sharing expertise and ensuring that work of this nature is possible wherever this disease occurs.

## Introduction

Colorectal cancer (CRC) is the second biggest cause of cancer-related deaths, globally. Although incidence has been higher in more developed countries, cases are rising in countries with traditionally lower rates (1). Over recent years, there has been a growing focus on the associations between the gut microbiome and the development of CRC, whether this is in the form of specific bacteria or bacterial toxins thought to have a role in carcinogenesis (2-4) or as an association with the overall gut flora, that could be informative as to the overall biology of the cancer, or useful in screening (5-7).

A major obstacle to a comprehensive understanding of the role of the microbiome in CRC biology (and microbiome studies in general) is the lack of diversity of sample populations. A recent meta-analysis of 444,829 publicly available microbiome samples from 2,592 studies found that, where origin was known, 71% of samples were from North America or Europe (8). The South Asian subcontinent was particularly under-represented, making up just 1.8% of samples, despite the region accounting for around a quarter of the global population, and an increasing CRC incidence (1). Studies focusing on Indian microbiomes have demonstrated a marked difference with Western populations (9-11), with taxa such as *Prevotella* being noticeably more abundant in Indian samples. This means that published associations between CRC and the microbiome may not be relevant to Indian (or other under-represented) patients. Studies of Indian populations have mainly sought to understand the microbiome of healthy individuals, or are group-specific studies comparing rural with urban, tribal or other localised population patterns. Sample numbers of studies examining the CRC microbiome in under-represented populations have mostly been limited, and have not been compared to global patterns, in order to better understand the similarities and differences between countries with different CRC incidences.

As part of an effort to address this imbalance, we have previously established an International CRC Microbiome Network seeking to advance the study of the microbiome of CRC in under-represented countries (namely India, Vietnam, Argentina and Chile in the initial phase). As an extension of the process of data generation, we participated in knowledge and expertise exchange. We also optimised cost-effective sample collection and storage protocols alongside sequencing and analysis strategies that are robust enough to be reproducible in countries which may not have the infrastructure of better-resourced facilities. We demonstrated that storing faecal samples on guaiac faecal occult blood (gFOBT) cards followed by 16S V4 rRNA sequencing allows long-term room-temperature storage and shipping of samples prior to processing, and produced data showing country specific microbial patterns as well as an international CRC bacterial signature (12).

This previous work was characterised by regional cancer centres in the aforementioned countries collecting samples, followed by shipping to the UK for processing. In the current study, we present the next logical step in the International CRC Microbiome Network aims of developing local expertise. We collected stool samples from a new cohort of 90 Indian CRC patients and healthy volunteers, and processed and sequenced them in India. We compared them to our previous data and to external cohorts to demonstrate the infrastructural and technical compatibility of the Indian institutions in carrying out microbial research into their own local cohorts, thereby demonstrating the value of country-specific analyses in globally significant fields such as CRC microbiome research.

## Methods

All research was carried out at The Cancer Institute (WIA), Chennai, India. Wherever possible, we used protocols developed during our previous work to establish a global CRC microbiome network (12), adapted for local use.

### Patient and healthy volunteer recruitment

Samples used in the study were collected between October 2018 and November 2019. Cases include CRC patients attending the Cancer Institute (WIA) and controls including healthy volunteers who were asymptomatic and unrelated to any gastrointestinal tract / genitourinary tract cancer patients.

### Ethical Approval

This study was performed with the approval from Institutional ethics committee (IEC/2018/01) at Cancer Institute (WIA), Chennai and Indian Council of Medical Research (2018-0337). All patients and healthy volunteers gave appropriate informed consent.

### Sample collection

Patients and healthy volunteers each provided a stool which was applied to all windows of a gFOBT card, and developer solution added within six hours. Once dry, cards were stored in sealed bags at ambient temperature until batch processing.

### DNA extraction

Three faecally loaded squares were excised using sterile scalpels, and DNA was extracted from gFOBT cards using a modified version of the QIAamp DNA mini kit (Qiagen, Germany) with additional Buffer ASL (Qiagen, Germany) as previously described (12).

### 16S rRNA library preparation and sequencing

We used the Earth Microbiome Project (EMP) 16S Illumina library preparation protocols (13) with Illumina 16S V4 primer constructs 515F (Parada)-806R (Apprill) (14), as previously described (12).

All libraries were pooled and sequenced by MedGenome Labs (Bangalore, India) on a single run of an Illumina MiSeq, using 2×250bp paired-end reads.

### Data processing

Reads were stripped of Illumina adapters using cutadapt (15). Further processing was carried out using QIIME2 (version 2019.10) (16). Reads were trimmed to 220bp to remove poor quality base calls, before denoising, pair merging and representative sequence assignment using DADA2 (17). Taxa were assigned to the SILVA database (version 132) (18) using BLAST+ (19), implemented within the QIIME2 q2 feature classifier plugin (20).

Within QIIME2, sequences were rarified to the level of the lowest-depth sample (43,000 QC-passing reads) before diversity analyses. Shannon index alpha diversity (21) of each sample was calculated, as well as Bray-Curtis beta diversity distances (22) between samples. Adonis PERMANOVA tests (23) were used to compare Bray-Curtis distances to sample metadata. Bray-Curtis distances were visualised using principle coordinate plots.

Taxa tables, and alpha and beta diversity measures were exported from QIIME2 for further analysis and visualisation in R (version 4.0.5). Random forest models (24) were built using randomForest (25) and validated using pROC (26). Taxa significantly associated with metadata were called using LEfSe (27).

### Comparisons to external data

We compared this dataset to our previous dataset (12) by merging QIIME2 tables, before processing as described above. We also compared to a metagenomic faecal dataset of Indian CRC samples (28), by examining genus-level taxonomic assignments of the two groups of samples.

## Results

### Sample numbers and sequencing quality

We recruited 47 CRC patients and 43 control healthy volunteers. Of these, one CRC patient failed sequencing QC, leaving 46 patients and 43 controls. Age and gender, comorbidities and information on the consumption of tobacco, meat and alcohol was collected for all participants, with additional, tumour-specific metadata available for the CRC patients, summarised in table 1.

**Table 1.** Patient and healthy volunteer demographics and metadata.

| **Sample details** | **Controls** | **CRC** |
| --- | --- | --- |
| **Total** | 43 | 46 |
| **Gender** |  |  |
| Male | 29 | 26 |
| Female | 24 | 20 |
| **Age** |  |  |
| Range | 23-76 | 21-76 |
| Median | 38 | 51 |
| **Comorbidities** |  |  |
| Diabetes | 6 | 9 |
| Hypertension | 1 | 6 |
| **Personal habits** |  |  |
| Smoker | 2 | 4 |
| Alcohol | 5 | 7 |
| Meat eater | 28 | 41 |
| **Cancer specific data** |  |  |
| **Site of cancer** | Right sided colon | 6 |
|  | Left sided colon | 15 |
|  | Rectum | 25 |
| **Grade** | Grade 1 | 1 |
|  | Grade 2 | 34 |
|  | Grade 3 | 11 |
| **Stage** | Stage 1 | 0 |
|  | Stage 2 | 8 |
|  | Stage 3 | 37 |
|  | Stage 4 | 1 |

Between 43,443 and 252,721 (median 120,789) sequences were called per sample, following pair merging and denoising by DADA2. This is a higher number than the Indian samples from our previous study (12), (minimum 58,984, maximum 128,124, median 106,324). However, feature number is mostly a reflection of sequencing depth. When all samples were rarefied to a similar depth and Shannon diversity compared (supplementary figure S1), the two datasets were comparable. The samples from the current study had a slightly higher diversity, but this was not significant (Mann-Whitney p=0.059). The slightly higher measured diversity could be a function of the sequencing technology used. The current study was performed using an Illumina MiSeq with 2×250bp reads to measure a 240bp PCR product, whilst the previous study used an Illumina HiSeq with 2×150bp reads. The MiSeq data deteriorated less over the span of the read, and combined with the longer read length might have made a range of representative sequences easier to reliably detect. Alternatively, the small sample numbers of the previous data (20 Indian samples) might have resulted in a slightly unrepresentative sample set.

Adonis PERMANOVA analysis of Bray-Curtis distances between the current study and the previous Indian samples, showed that sample status (cancer vs healthy volunteer) was associated with the largest proportion of measured differences, followed by which dataset the sample belonged, then gender and age (supplementary figure S2).

### Comparison with previous data

We first compared the current data with our previous global comparison using principle coordinate plots of Bray-Curtis distances (Fig 1). As we saw with our previous study, there appeared to be more geographic separation than separation by cancer status. Although the plot is now dominated by Indian samples, so biased, the South American samples are all in the left side of the plot, and Vietnamese samples mostly in the top half. Adonis PERMANOVA analysis indicated that sequencing run was associated with 3.4% of variation, country of origin with 6.7%, and cancer status with 2.3%, all of which were significant (p < 0.001).

**Figure 1.**
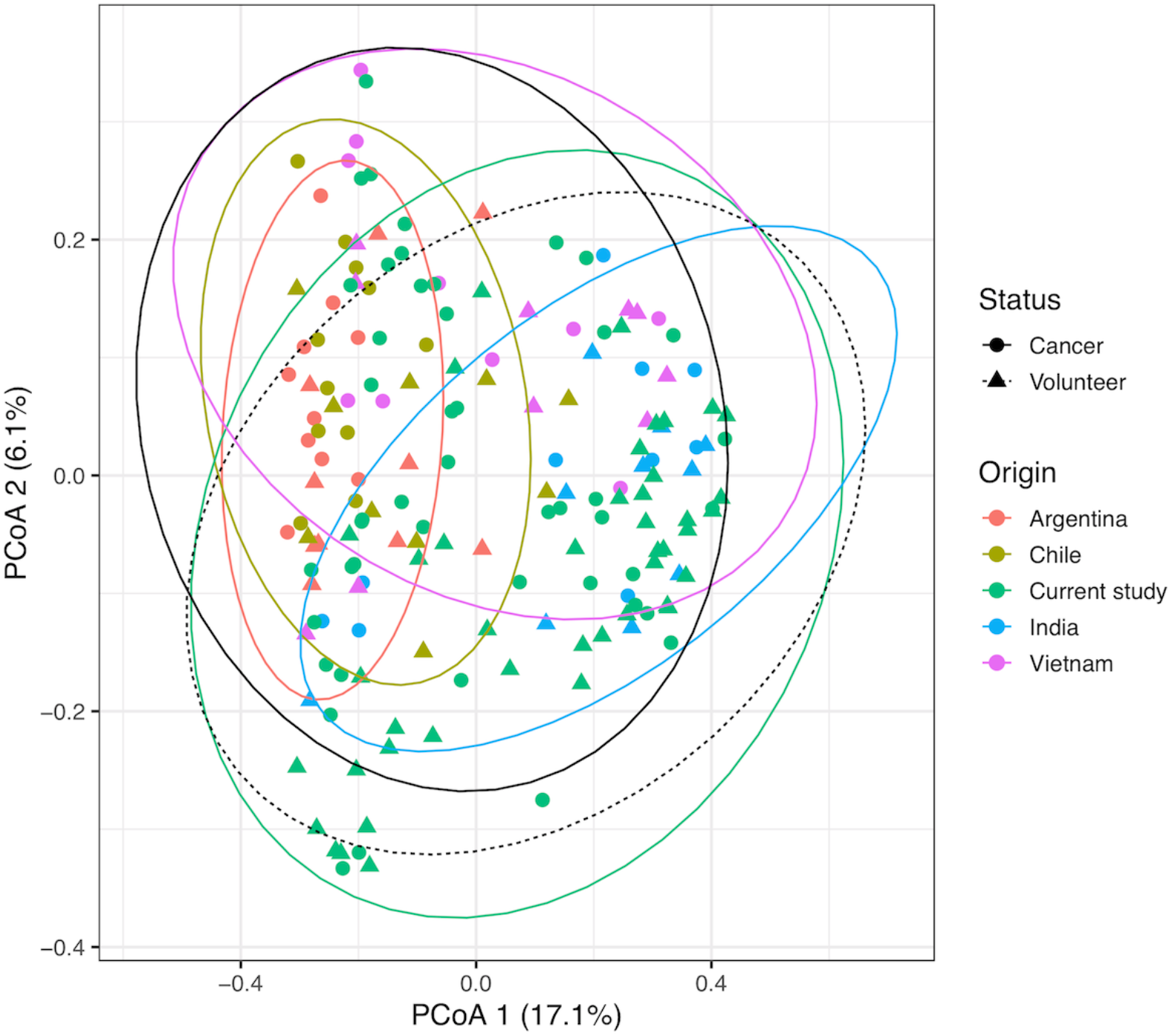
Principle coordinate plot of Bray-Curtis distances of the current study, compared to global samples from Young *et al*. “India” refers to previous Indian samples. 95% confidence intervals for sample groupings are displayed as ellipses.

Next, we compared the most prevalent taxa of the two datasets (Fig 2). As with the principle coordinate plot, there was clear geographical separation. The Asian cohorts were characterised by far greater abundance of *Prevotella*, and relatively less *Bacteroides*. All the cancer cohorts had more *Escherichia/Shigella* than their respective control cohorts. We tested for taxa differentiating between cancer and healthy volunteer for a merged dataset of all countries from both cohorts, and again with just Indian samples, using LEfSe (27). This allowed us to examine taxa outside of the most common genera. (Supplementary figure S3). For both comparisons, known CRC taxa such as *Akkermansia, Fusobacterium* and *Ruminococcus* were evident, reiterating our previous findings that there appears to be a consistent CRC-associated faecal flora.

**Figure 2.**
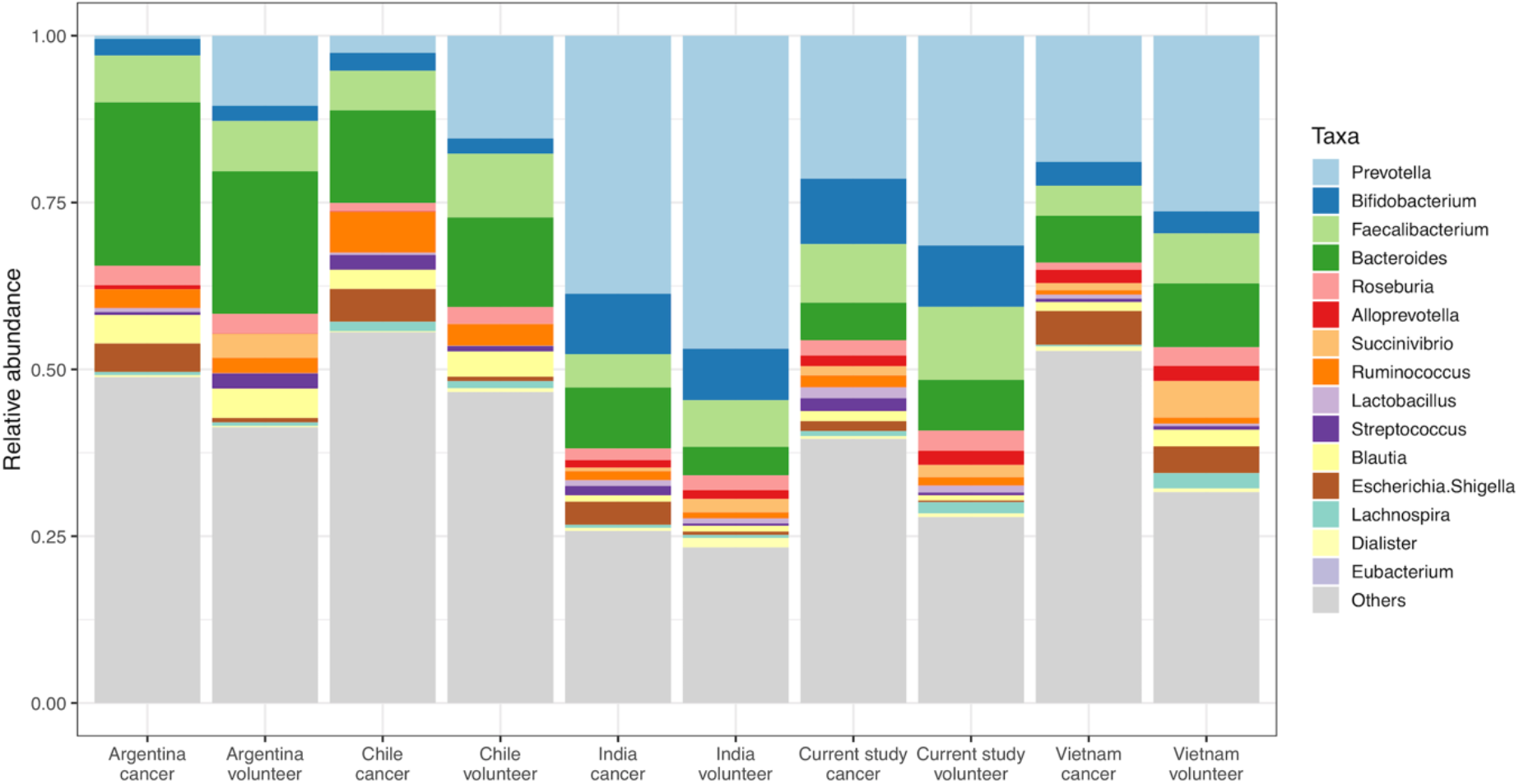
Taxa abundance of the current study and Young *et al*. The top 15 genera across all cohorts are displayed.

We also compared alpha diversity of cancer versus healthy volunteer for the different countries (figure 3). As we had previously showed, Indian (and Vietnamese) samples have lower diversity than South American samples. It was previously seen that cancer samples had higher diversity than healthy volunteers for all the countries, although the small sample numbers for the respective countries made this difficult to claim with certainty. This trend can now be confirmed for Indian samples, given the increased sample number (Mann-Whitney p=0.0005).

**Figure 3.**
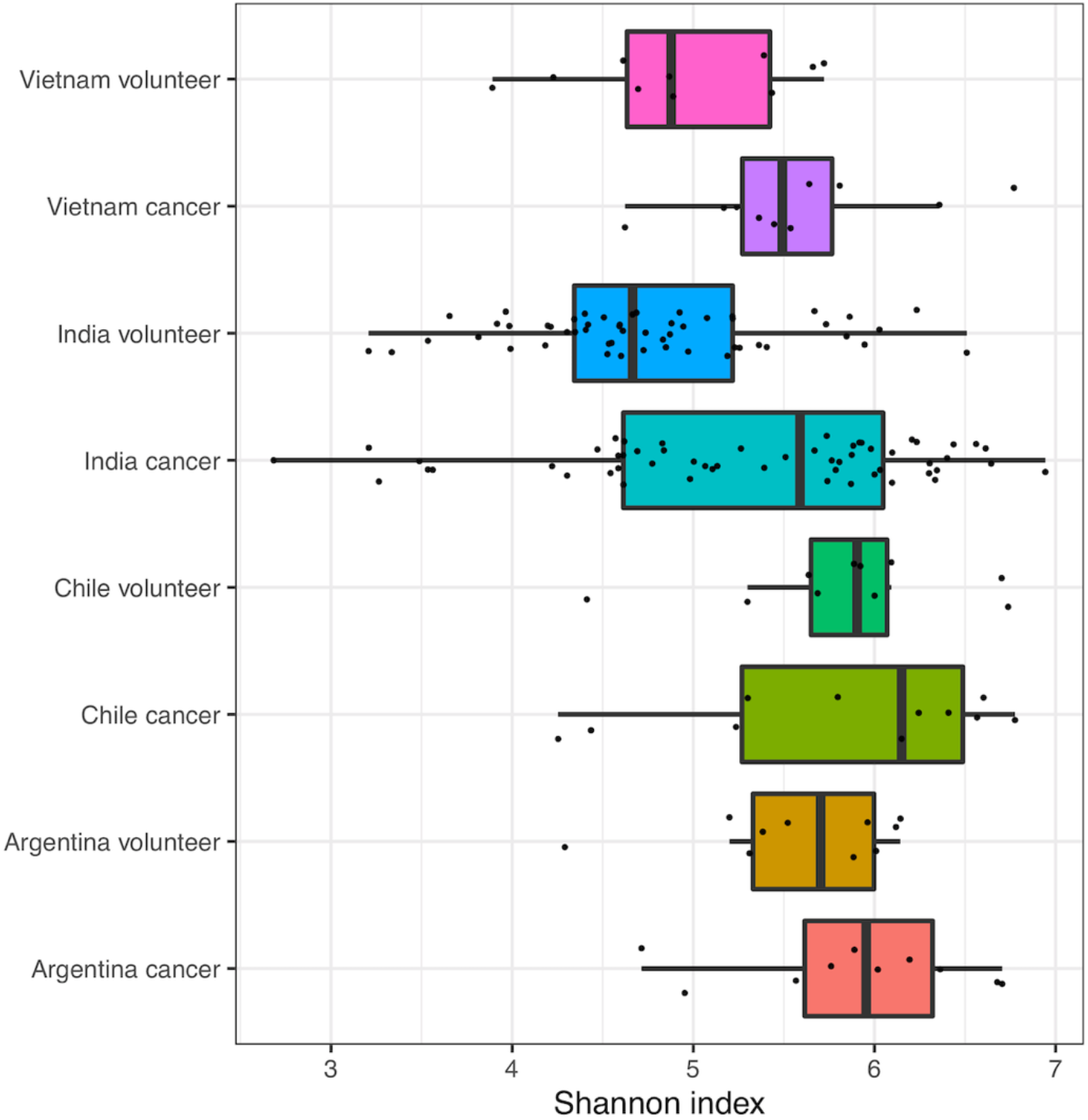
Shannon index alpha diversity of the current study and Young *et al*. Indian samples from the two studies have been merged.

### Prediction of cancer status using previous data

As well as visually inspecting similarities between our previous global dataset and our current study, we sought to ascertain if the previous data could be used to predict cancer status in the current samples, and vice versa.

For both the Young *et al*. cohort and our new cohort, we generated random forest models and used them to predict cancer status in their own samples, and then validated them in the other dataset (Figure 4). There was very good concordance. The previous samples had an area under the curve (AUC) of 0.77 when used to predict their own samples, which only dropped to 0.76 when our current study was used as the original model. Our current study had an internal AUC of 0.86, which dropped to 0.85 when cancer status was predicted using the model from the Young *et al*. cohort. It is interesting that prediction status of our current dataset based on the model of our previous samples performs better than our previous cohort predicting the status of its own samples. Perhaps using samples from only one country gives a more homogenous dataset, or maybe Indian samples are easier to predict, with more dramatic differences in levels of predictive taxa such as *Prevotella* between cancer patients and healthy volunteers. The theory that Indian samples may be easier to predict is corroborated by meta-analysis carried out within the Young *et al*. paper. Ten studies, including that of Young *et al*. and the Gupta *et al*. cohort if Indian CRC and healthy volunteers. A random forest model was built of each cohort and validated against every other cohort. The validation using the Indian Gupta cohort had the highest AUC value for five of the nine models (including the model based on Young *et al*.) as well as the highest mean AUC.

**Figure 4:**
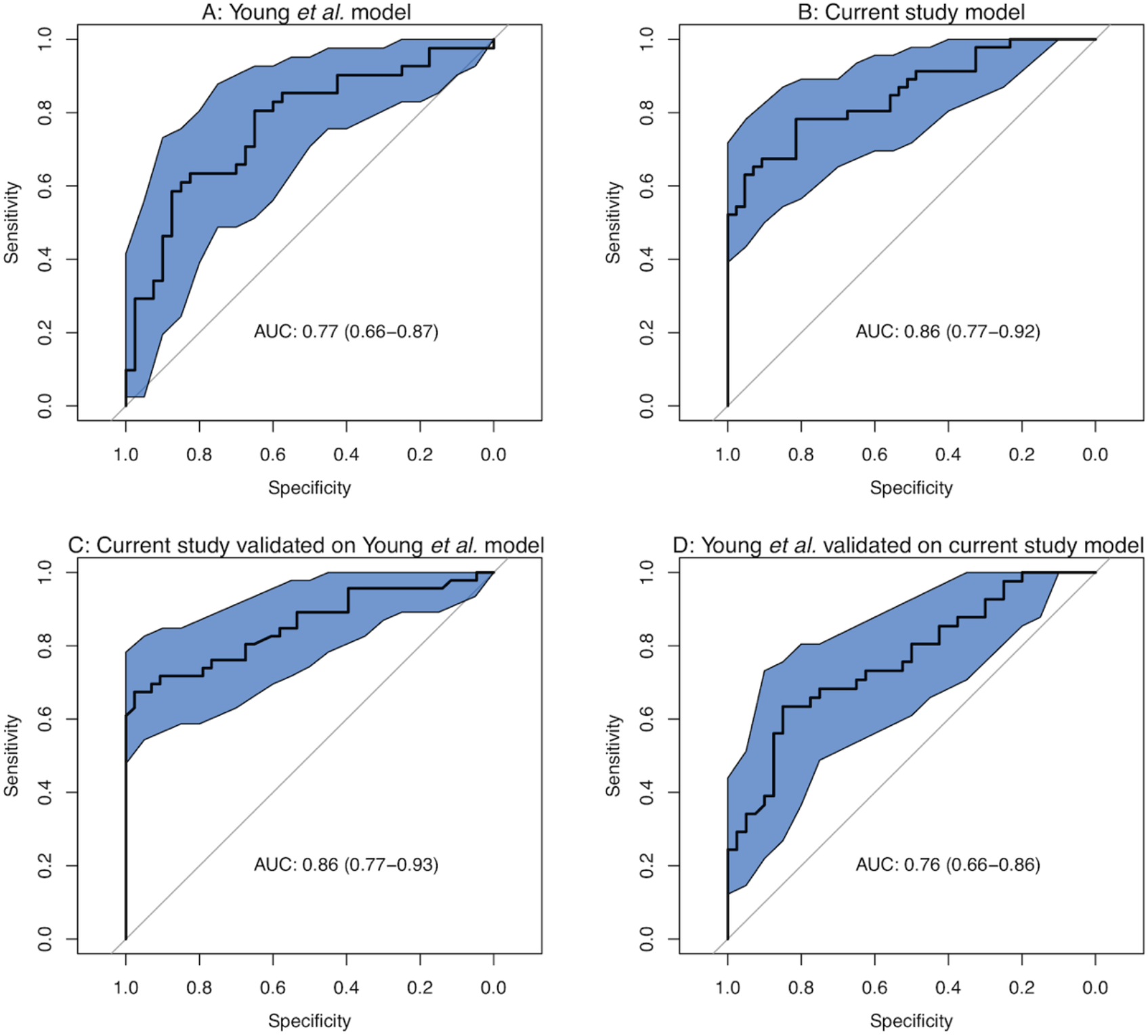
Using random forest models of (A) our previous global dataset and (B) our current study to predict cancer status in their own samples and (C, D) to predict cancer status in samples from the other study.

These results repeat our previous findings that there appears to be a consistent, global pattern of changes in the faecal microbiome of CRC patients, and that if this is adequately catalogued, can be used to predict the cancer status of new samples.

### Comparisons with other Indian datasets

There are a limited number of studies profiling Indian faecal microbiomes. We visually compared our data to three studies profiling healthy Indian healthy volunteers (9-11). Whilst these studies had different aims, largely based on profiling the variation across India and between Indian and non-Indian populations, they were all characterised by the dominance of *Prevotella* amongst a large proportion of their samples, as we observed in both our healthy volunteers and cancer patients.

We were only aware of one other study comparing the microbiomes of CRC patients to healthy volunteers in Indian samples, based on individuals from Bhopal and Kerala (28). We converted the species calls of the Gupta metagenomic dataset into genus calls, to match our 16S genus calls, then compared the CRC and healthy volunteer taxa in both groups of samples.

Visually, whilst there were differences, some patterns were consistent between the datasets (Figure 5). *Prevotella* was higher in control populations for both groups while *Escherichia/Shigella* was higher in CRC samples. When comparing using LEfSe (Supplementary figure S3), there were several similarities. Out of 43 CRC-associated taxa in the current study, 13 were amongst the 28 CRC-associated taxa from Gupta *et al*. Out of control-associated taxa, two were shared between four taxa called from the current study and six from Gupta *et al*. The main discrepancy was that *Bacteroides* was associated with healthy controls in the current study and CRC in the Gupta cohort, possibly due to differences between 16S and whole metagenomic sequencing strategies, but more likely due to using different cohorts from different regions of India. Different populations may have different prevalent species of *Bacteroides*, with some relatively benign and some containing CRC-associated strains (29).

**Figure 5:**
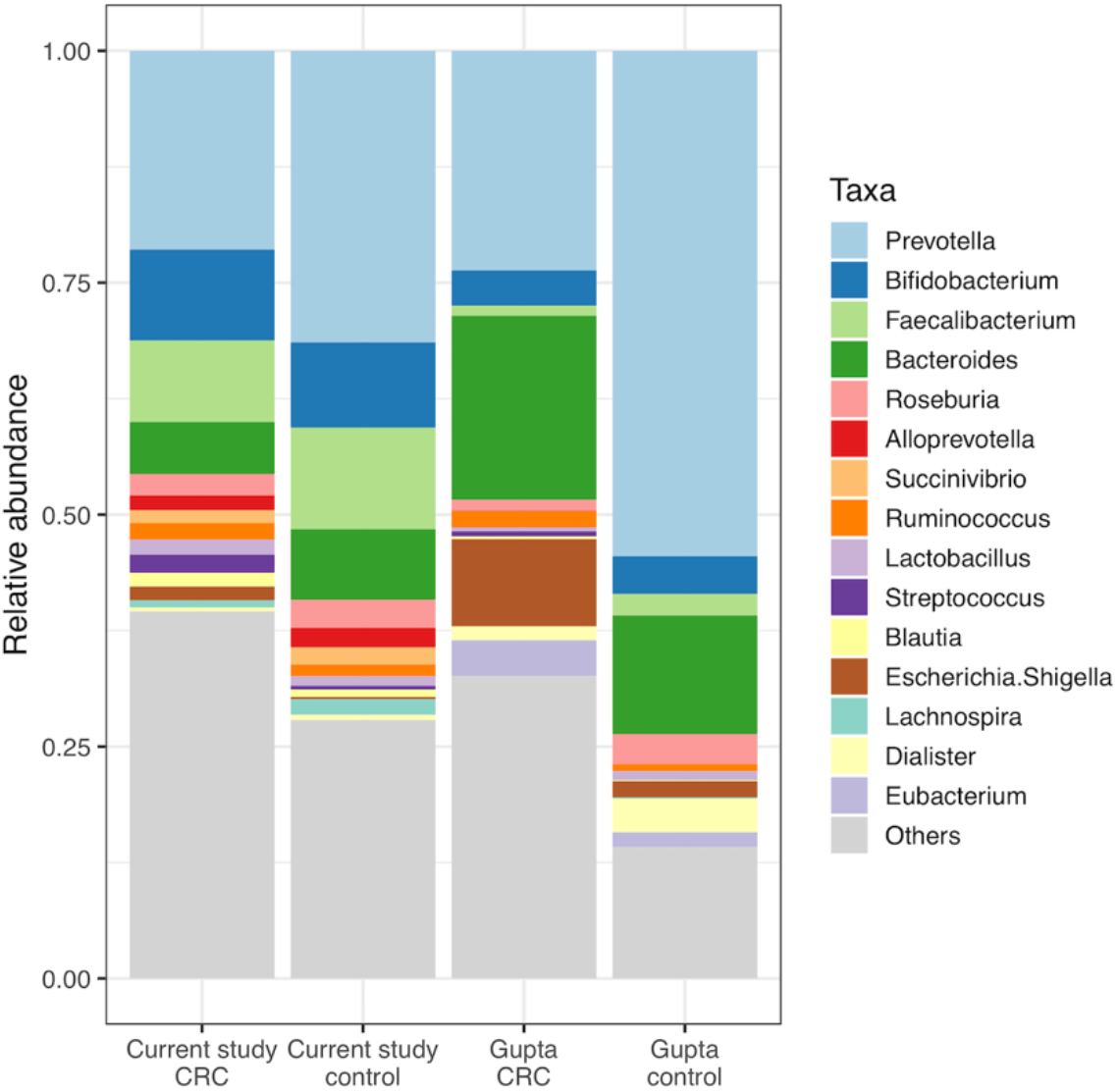
Taxa abundance of the current study and Gupta *et al*. The top 15 genera across all cohorts are displayed.

Adonis PERMANOVA analysis of location and cancer status of the merged datasets suggested that location within India was associated with 18% of the variation, whilst cancer status was associated with 4.4%. However, it is impossible to know how much of the apparent location-associated variation is in fact a product of sequencing strategy. By removing our samples, thus keeping sequencing strategy uniform, location was associated with 7.8% of variation and cancer status with 10.8%. All associations in both comparisons were significant (p < 0.001).

As well as simply counting taxa which fell into different subgroups, we built random forest models of our data and that of Gupta *et al*. and validated them with the other dataset (Figure 6). Again, there was good concordance. The cancer status of both datasets were predicted better using their own samples than the models built with the alternative dataset, but with overlapping confidence intervals. This demonstrates again that taxa calls from one dataset can reliably predict cancer status in another, despite completely different sequencing strategies being used.

**Figure 6:**
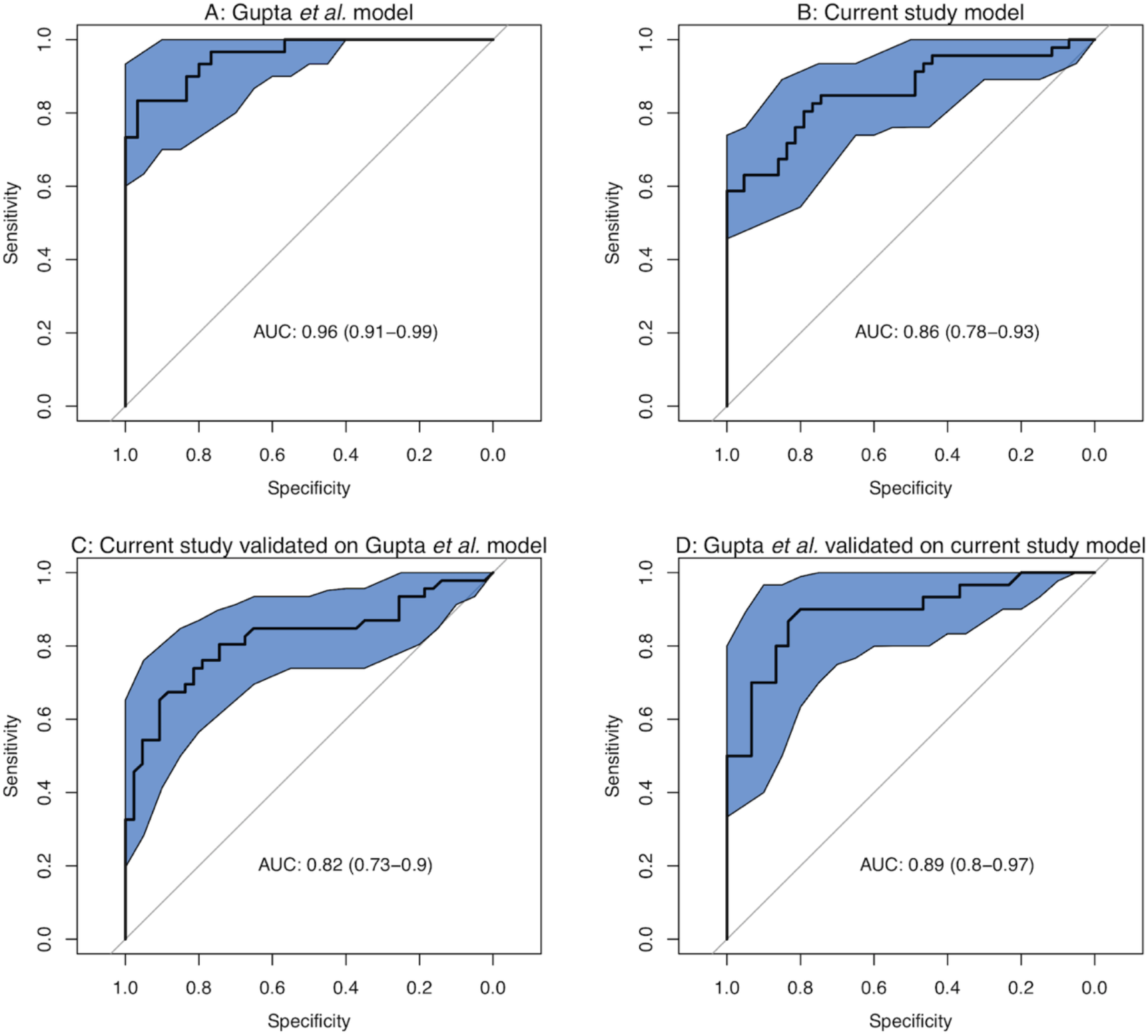
Using random forest models of (A) Gupta *et al*. and (B) our current study to predict cancer status in samples in their own samples and (C,D) to predict cancer status in samples from the other study.

### Associations with metadata

Lastly, and acknowledging the relatively small size of our cohort, we examined whether any of the different patient groupings of our CRC samples were associated with changes in the microbiome.

When comparing different metadata categories using adonis PERMANOVA of Bray-Curtis distances, only anatomical site and sex were significant. We compared anatomy, age and sex on a principle coordinate plot which reiterated these findings. Although there was overlap, all the right-sided colon samples were in the top half of the plot, and almost all the female samples in the left half (Figure 7).

**Figure 7:**
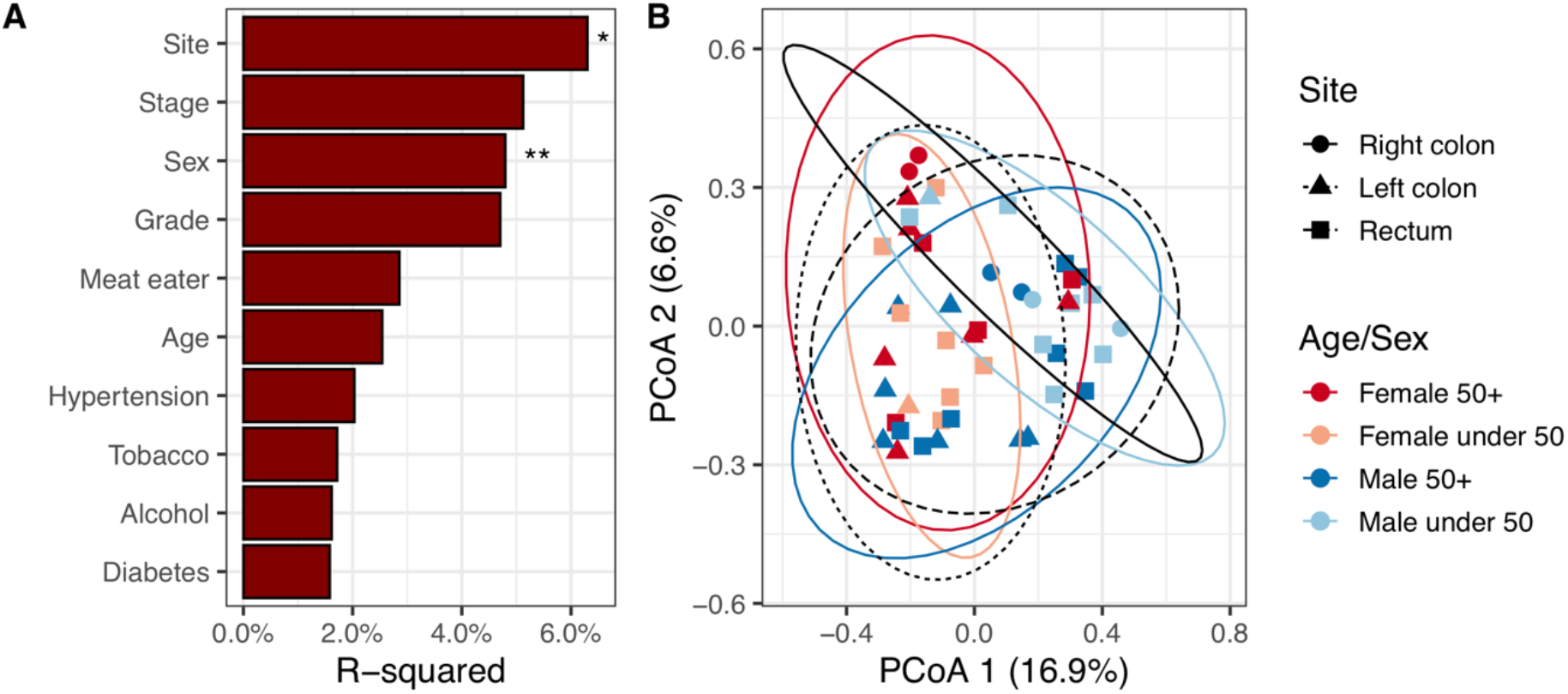
Beta diversity analyses of current study cancer samples. (A) Adonis PERMANOVA comparison with metadata. R-squared refers to amount of Bray-Curtis variation associated with each category. P-value is indicated by: ** - p <= 0.01; * - p <= 0.05. (B) Principle coordinate plot of Bray-Curtis distances. Shape of points refers to anatomical site, while colour refers to age and sex. 95% confidence intervals for sample groupings are displayed as ellipses.

When comparing taxa for the different metadata categories (Figure 8), a number of differences could be seen, most noticeably in the relative abundances of the most common genera *Prevotella, Bifidobacterium, Faecalibacterium* and *Bacteroides*. It should be noted that some sample numbers of some groupings are very small. For instance, there were only six right-sided colon samples. Therefore, we are unwilling to analyse this aspect of the data too deeply, and present it more as an example of what kind of analyses would be possible with larger sample numbers.

**Figure 8:**
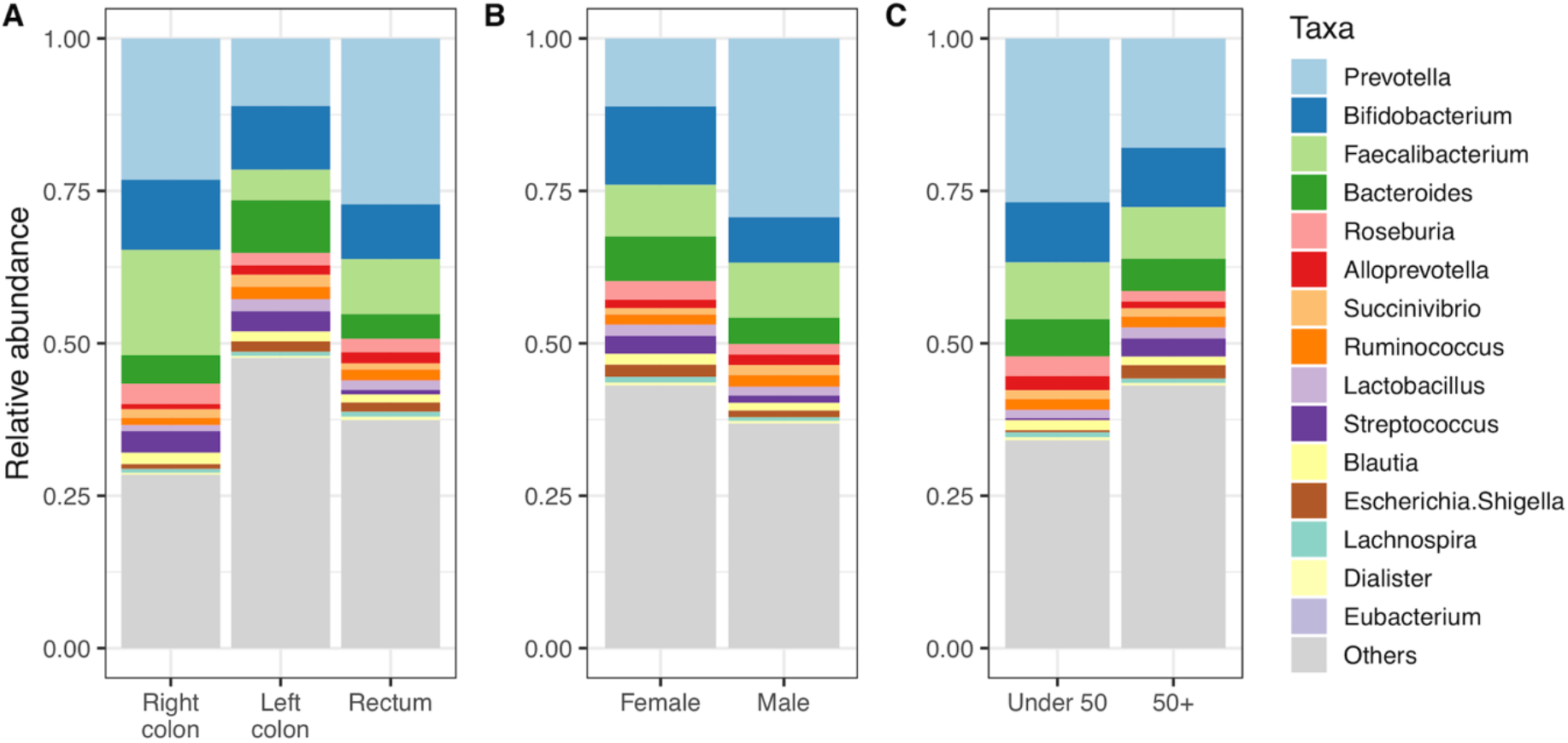
Taxa abundances of the cancer samples split by (A) anatomical site, (B) sex and (C) age.

## Discussion

Given the known paucity of microbial datasets from outside of Europe and North America (8), it is vital for the community to strive to increase the number and size of available datasets for a better understanding of the role of the microbiome in health and disease. In particular, it is important to gain an understanding of which groups of taxa or specific components of the microbiome such as toxins (4) are important for non-invasive CRC diagnosis, or as potential therapeutic targets. As the field of CRC microbiome studies advances, it is important to know which developments are universal, and which are confined to limited geographical regions. This is particularly evident when examining the limited number of Indian microbial datasets (9-11, 28) which consistently report that *Prevotella* is the dominant genus present, in contrast with most other countries examined. In this context, it becomes important to know if putative oncomicrobes or potential screening targets are of value in Indian populations.

Having previously set up an international network of institutions from countries with under-developed microbial research infrastructure, we have been developing robust protocols and producing CRC microbiome data from those countries, demonstrating marked differences between regions, yet with promising evidence of a universal CRC faecal microbiome (12). As a critical output resulting from the establishment of that knowledge transfer network, we are now able to present a new cohort of CRC and control microbiomes, wholly collected and processed in India. We compared these samples to the Indian samples from our previous collection, sequenced in the UK, and found them to be largely comparable. The new data was of slightly higher quality, and separated from the old data in diversity analyses. We attribute this to the sequencing technology used, with less quality deterioration across the length of sequencing reads observed in the MiSeq reads than in the HiSeq reads of the previous study. Therefore, we have demonstrated that Indian institutions can advance their microbiome research associated with healthcare benefits, in a systematic manner comparable with that of the datasets produced elsewhere.

Moving from logistics to biology, we compared our new dataset to all the samples from our international network and external studies. As seen before, our Indian samples were characterised by a high abundance of *Prevotella*, especially in the healthy volunteers, and an increase in alpha diversity amongst the cancer samples. It has previously been suggested that the dominance of *Prevotella* in healthy Indian samples leads to low diversity, and that the relative decrease in *Prevotella* in Indian CRC samples allows more taxa to flourish, resulting in increased diversity (28).

There were differences between our dataset and that of Gupta *et al*. Despite being the dominant genus, *Prevotella* was less abundant in our samples. This could be a bias of sequencing strategy, or it could be differences between Indian regions. Comparisons of the datasets with and without mixed sequencing strategies suggested that both factors had measurable effects. Our samples were all from Chennai region, whereas the Gupta samples were taken from Bhopal and Kerala, several hundred kilometres away, with different local conditions and prevalent diets. It needs to be highlighted at this juncture that the socio-economic differences, and regional/cultural diversity will have a major effect on the dietary patterns prevalent in Indian studies and will influence the health status of individuals. Even within India, these differences will affect microbial patterns associated with the gut microbiome in general, and CRC in particular. Previous comparisons of healthy Indian samples have shown considerable regional variation in *Prevotella* and other genera, thought to be largely due to diet, specifically levels of meat intake (9-11). It is not possible to make exact comparisons with diet between our study and the Gupta cohort, which has no dietary information recorded. A historical study across India (30) suggested that Chennai (21%) has levels of vegetarianism intermediate to that of Bhopal (45%) and Kerala (6%). 22% of our cohort were vegetarians, although the small sample size means that we cannot infer that this is reflective of the population as a whole.

When we used models from our previous and new datasets to predict the cancer status of each other, there was good concordance. Only slight drops of predictive success were observed between internal and external validation. This pattern was repeated when we compared our new dataset with that of Gupta *et al*., despite the Gupta dataset using a completely different sequencing strategy (metagenomic shotgun sequencing compared to 16S rRNA amplicon sequencing, each of which introduce different biases). Together, these comparisons show that while there does seem to be a universal CRC-microbiome, and that findings can be generalised across continents, there are more similarities between datasets collected within countries, and that efforts should continue to improve the number of samples available from under-represented regions. It may well be that using local control datasets may allow us to improve the predictive value of microbiome studies in bowel cancer screening.

## Data availability

Raw sequence data is available from the European Nucleotide Archive, accession number PRJEB53415 (http://www.ebi.ac.uk/ena/data/view/PRJEB53415).

A STORMS (Strengthening The Organizing and Reporting of Microbiome Studies) (31) checklist 1.03 is available at doi:10.5281/zenodo.6839159

## Supplementary material

Table S1 contains genus-level taxonomic calls for each sample in this study.

Figures S1-S3 contain alpha and beta diversity statistics, plus taxa associating with CRC in various combinations of samples.

