## Supplementary figures for "Analysis of an Indian colorectal cancer faecal microbiome collection demonstrates universal colorectal cancer-associated patterns, but closest correlation with other Indian cohorts"

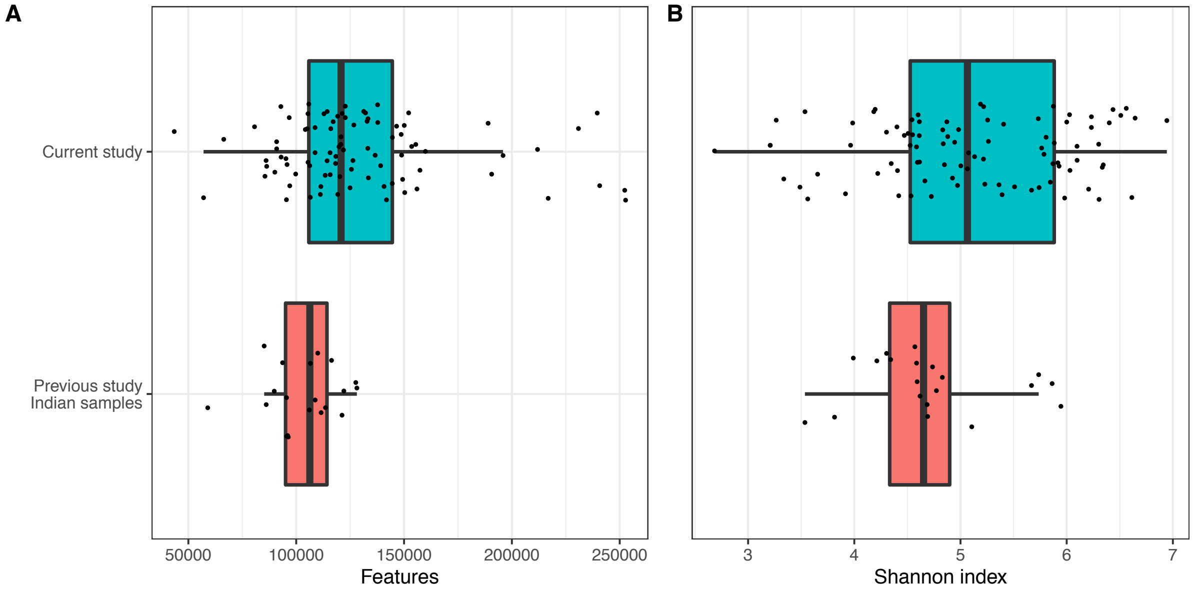


Supplementary figure S1: Number of features per sample called by DADA2 (A) and Shannon index alpha diversity (B) for the current study and the Indian samples from our previous study, Young *et al*.

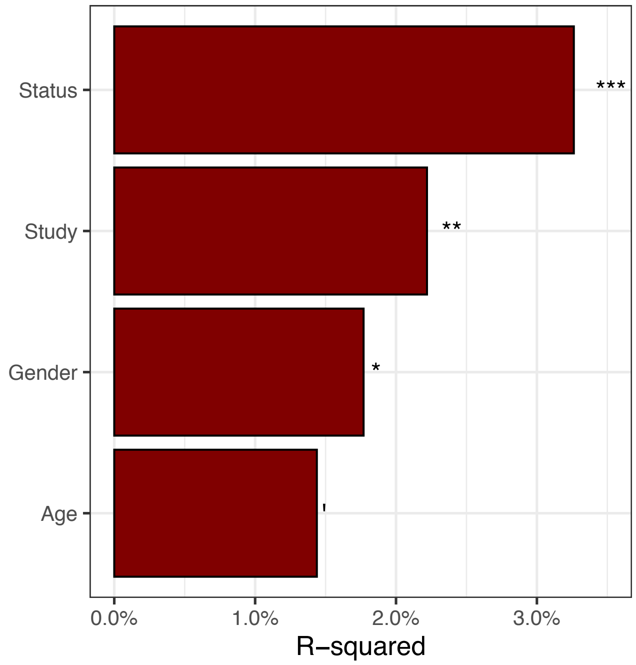


Supplementary figure S2. Adonis PERMANOVA comparison of the current study and Indian samples from our previous study. R-squared refers to amount of Bray-Curtis variation associated with each metadata category. Status is cancer vs healthy volunteer. Study refers to the current study vs the previous work. P-value is indicated by: *** - p <= 0.001; ** - p <= 0.01; * - p <= 0.05; ‘ – p <= 0.1.

Supplementary figure S3 (next four pages). LEfSe results comparing (A) cancer versus volunteer for merged datasets of Young *et al* with the current study, (B) just the Indian samples of Young *et al* and the current study, (C) just the current study, and (D) metagenomic samples from Gupta *et al.*

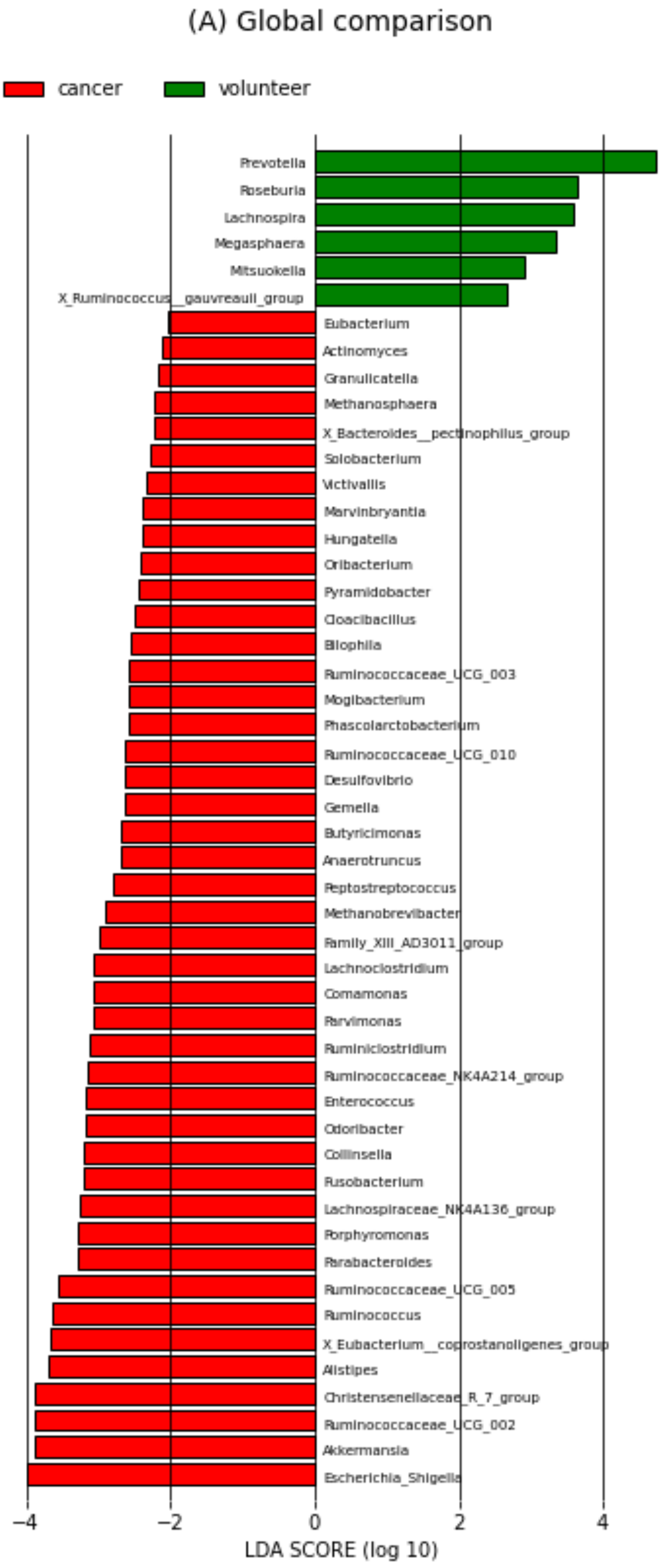
(A) Global comparison
cancer
volunteer
Prevotella
Roseburia
Lachnospira
Megasphaera
Mitsuokella
X_Ruminococcus__gauvreauii_group
Eubacterium
Actinomyces
Granulicatella
Methanosphaera
X_Bacteroides__pectinophilus_group
Solobacterium
Victivallis
Marvinbryantia
Hungatella
Oribacterium
Pyramidobacter
Cloacibacillus
Bilophila
Ruminococcaceae_UCG_003
Mogibacterium
Phascolarctobacterium
Ruminococcaceae_UCG_010
Desulfovibrio
Gemella
Butyricimonas
Anaerotruncus
Peptostreptococcus
Methanobrevibacter
Family_XIII_AD3011_group
Lachnoclostridium
Comamonas
Parvimonas
Ruminiclostridium
Ruminococcaceae_NK4A214_group
Enterococcus
Odoribacter
Collinsella
Fusobacterium
Lachnospiraceae_NK4A136_group
Porphyromonas
Parabacteroides
Ruminococcaceae_UCG_005
Ruminococcus
X_Eubacterium__coprostanoligenes_group
Alistipes
Christensenellaceae_R_7_group
Ruminococcaceae_UCG_002
Akkermansia
Escherichia_Shigella
−4
−2
0
2
4
LDA SCORE (log 10)

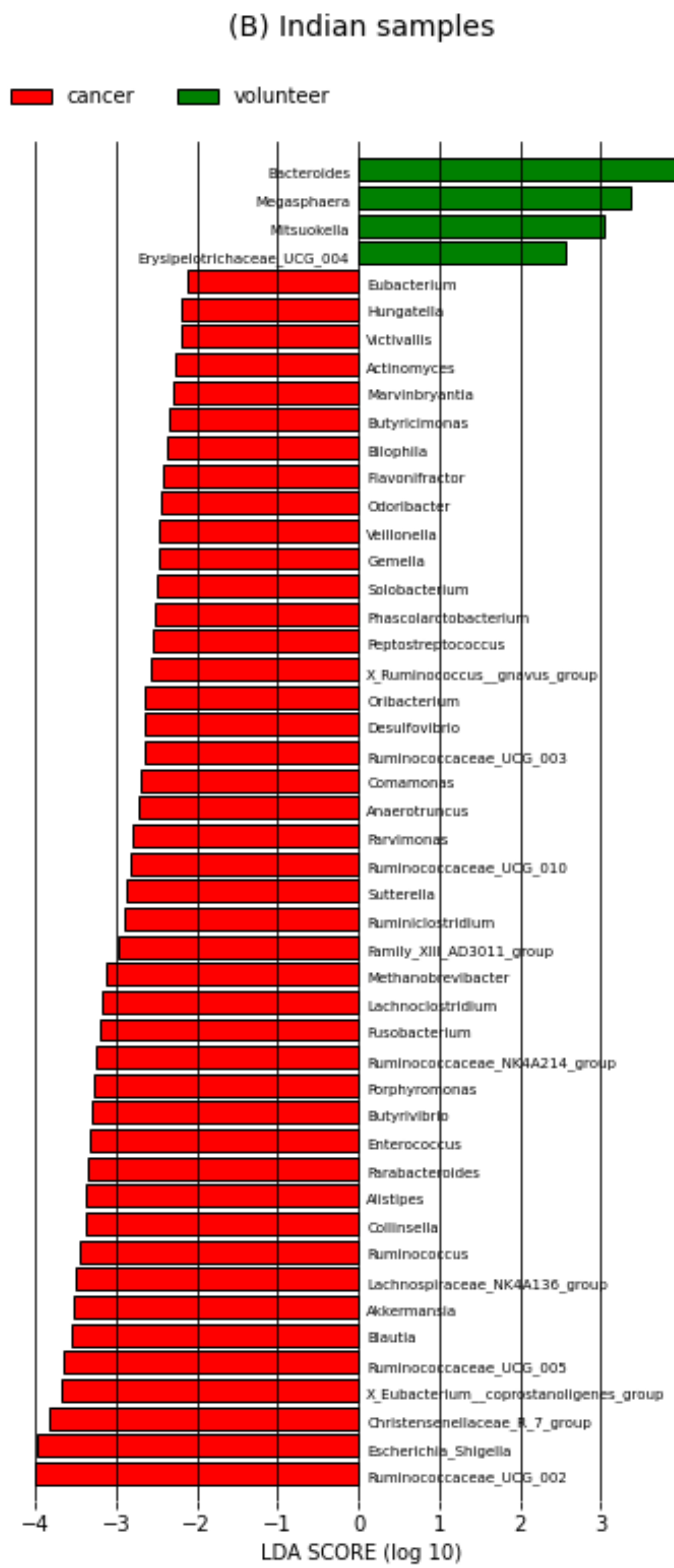
(B) Indian samples
cancer
volunteer
Bacteroides
Megasphaera
Mitsuokella
Erysipelotrichaceae_UCG_004
Eubacterium
Hungatella
Victivallis
Actinomyces
Marvinbryantia
Butyricimonas
Bilophila
Flavonifractor
Odoribacter
Veillonella
Gemella
Solobacterium
Phascolarctobacterium
Peptostreptococcus
X_Ruminococcus__gnavus_group
Oribacterium
Desulfovibrio
Ruminococcaceae_UCG_003
Comamonas
Anaerotruncus
Parvimonas
Ruminococcaceae_UCG_010
Sutterella
Ruminiclostridium
Family_XIII_AD3011_group
Methanobrevibacter
Lachnoclostridium
Fusobacterium
Ruminococcaceae_NK4A214_group
Porphyromonas
Butyrivibrio
Enterococcus
Parabacteroides
Alistipes
Collinsella
Ruminococcus
Lachnospiraceae_NK4A136_group
Akkermansia
Blautia
Ruminococcaceae_UCG_005
X_Eubacterium__coprostanoligenes_group
Christensenellaceae_R_7_group
Escherichia_Shigella
Ruminococcaceae_UCG_002
−4
−3
−2
−1
0
1
2
3
4
LDA SCORE (log 10)

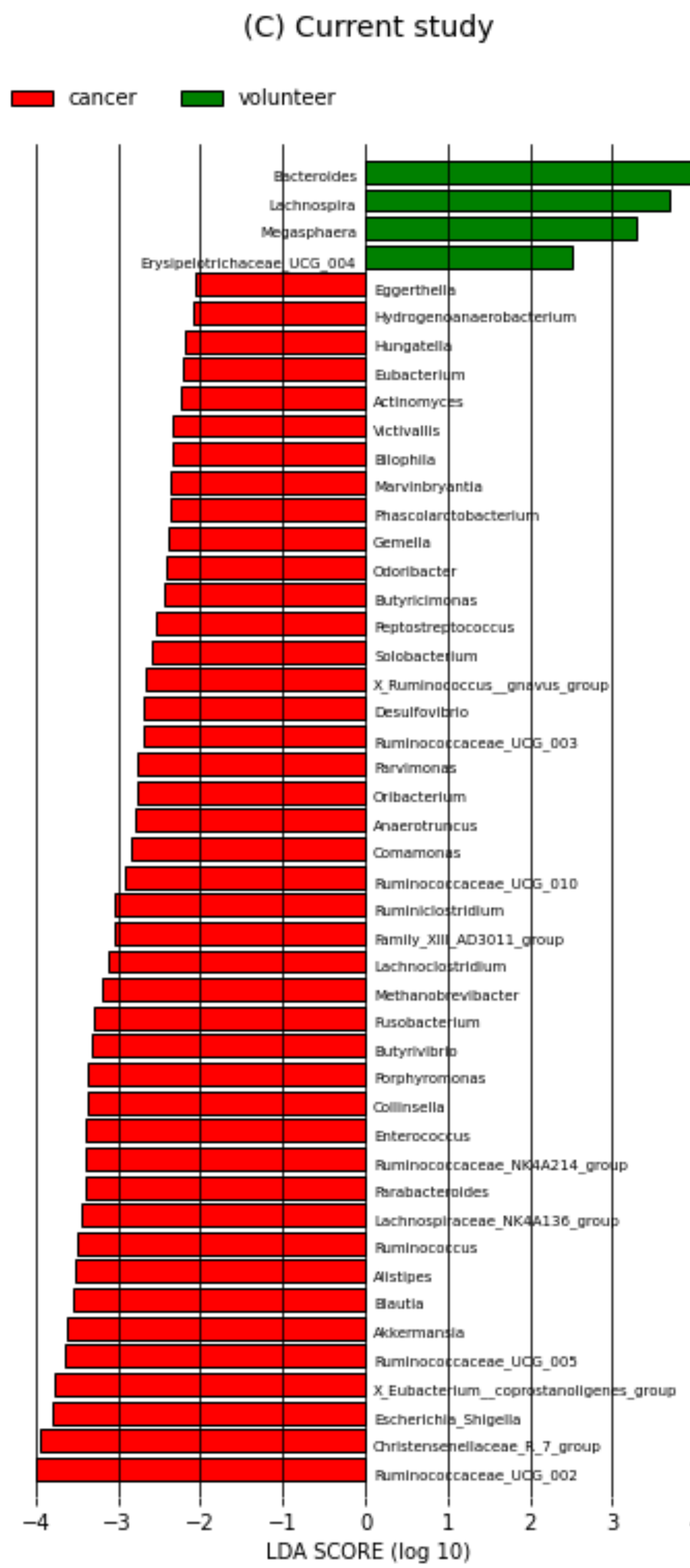

(C) Current study
cancer
volunteer
Bacteroides
Lachnospira
Megasphaera
Erysipelotrichaceae_UCG_004
Eggerthella
Hydrogenoanaerobacterium
Hungatella
Eubacterium
Actinomyces
Victivallis
Bilophila
Marvinbryantia
Phascolarctobacterium
Gemella
Odoribacter
Butyricimonas
Peptostreptococcus
Solobacterium
X_Ruminococcus__gnavus_group
Desulfovibrio
Ruminococcaceae_UCG_003
Parvimonas
Oribacterium
Anaerotruncus
Comamonas
Ruminococcaceae_UCG_010
Ruminiclostridium
Family_XIII_AD3011_group
Lachnoclostridium
Methanobrevibacter
Fusobacterium
Butyrivibrio
Porphyromonas
Collinsella
Enterococcus
Ruminococcaceae_NK4A214_group
Parabacteroides
Lachnospiraceae_NK4A136_group
Ruminococcus
Alistipes
Blautia
Akkermansia
Ruminococcaceae_UCG_005
X_Eubacterium__coprostanoligenes_group
Escherichia_Shigella
Christensenellaceae_R_7_group
Ruminococcaceae_UCG_002
−4
−3
−2
−1
0
1
2
3
4
LDA SCORE (log 10)

(D) Gupta et al.

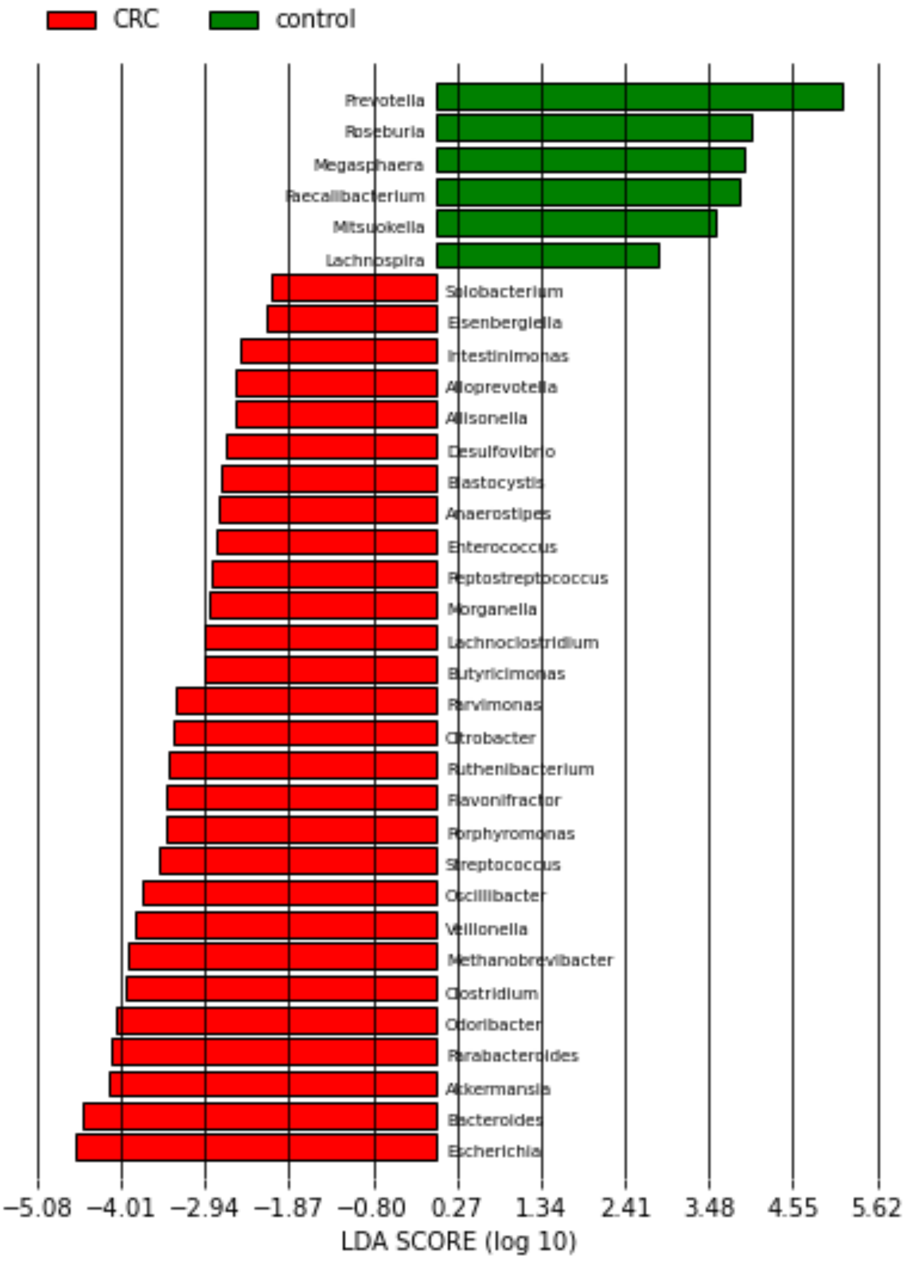
CRC
control
Prevotella
Roseburia
Megasphaera
Faecalibacterium
Mitsuokella
Lachnospira
Solobacterium
Eisenbergiella
Intestinimonas
Alloprevotella
Allisonella
Desulfovibrio
Blastocystis
Anaerostipes
Enterococcus
Peptostreptococcus
Morganella
Lachnoclostridium
Butyricimonas
Parvimonas
Citrobacter
Ruthenibacterium
Flavonifractor
Porphyromonas
Streptococcus
Oscillibacter
Veillonella
Methanobrevibacter
Clostridium
Odoribacter
Parabacteroides
Akkermansia
Bacteroides
Escherichia
−5.08
−4.01
−2.94
−1.87
−0.80
0.27
1.34
2.41
3.48
4.55
5.62
LDA SCORE (log 10)
